# Can deep convolutional neural networks support relational reasoning in the same-different task?

**DOI:** 10.1101/2021.09.03.458919

**Authors:** Guillermo Puebla, Jeffrey S. Bowers

## Abstract

Same-different visual reasoning is a basic skill central to abstract combinatorial thought. This fact has lead neural networks researchers to test same-different classification on deep convolutional neural networks (DCNNs), which has resulted in a controversy regarding whether this skill is within the capacity of these models. However, most tests of same-different classification rely on testing on images that come from the same pixel-level distribution as the testing images, yielding the results inconclusive. In this study we tested relational same-different reasoning DCNNs. In a series of simulations we show that models based on the `ResNet-50` architecture are capable of visual same-different classification, but only when the test images are similar to the training images at the pixel-level. In contrast, even when there are only subtle differences between the testing and training images, the performance of DCNNs drops substantially. This is true even when DCNNs’ training regime is augmented with images from new versions of the same-different task or through multi-task learning on the test images. Furthermore, we show that the Relation Network, a deep learning architecture specifically designed to tackle visual relational reasoning problems, suffers the same kind of limitations than `ResNet-50` classifiers.

## 1 Introduction

Relational reasoning is core to human intelligence (Penn, Holyoak, & Povinelli, 2008), and has proven to be a challenge for an earlier generation of connectionist models (e.g., O’Reilly & Busby, 2002; Rogers & McClelland, 2004; St. John, 1992) as well as more recent deep neural networks (for recent reviews, see Ricci, Cadène, & Serre, 2021; Stabinger, David, Piater, & Rodríguez-Sánchez, 2020). Perhaps the simplest form of relational reasoning is the same-different task that simply requires the reasoner to determine whether two inputs are the same or different by some criterion. In the domain of vision, the simplest version of this is to classify images as visually identical or not. This skill, essential to abstract combinatorial thought, is much more developed in humans and chimpanzees than in other species (Gentner, Shao, Simms, & Hespos, 2021), and develops early in human infants (e.g., Ferry, Hespos, & Gentner, 2015).

Recently there has been mixed evidence regarding whether standard deep convolutional neural networks (DCNNs) can support same-different matching of images. Much of this research has used the Synthetic Visual Reasoning Test developed by Fleuret et al. (2011). This dataset comprises set of 23 classification problems involving images of randomly generated shapes (for example images see Fig. 2A). In his study Fleuret et al. (2011) found that the standard machine learning techniques of the time performed poorly while most humans were able to solve the problems after seeing a few examples. Similarly, Stabinger, Rodríguez-Sánchez, and Piater (2016) showed that state-of-the-art DCNNs (at the time) LeNet and GoogLeNet performed poorly on the same SVRT same-different tasks, and more recently, Kim et al. (2018) showed that vanilla DCNNs were poor at SVRT same-different tasks, and using a different dataset, showed that Santoro et al. (2017) relational network (RN) also failed to support same-different judgments.

**Figure 1:**
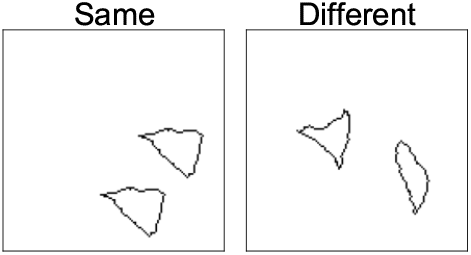
Examples of the “same” and “different” categories from SVRT problem #1. In this problem an image belongs to the category “same” if both shapes are identical up to translation on the canvas and “different” otherwise.

**Figure 2:**
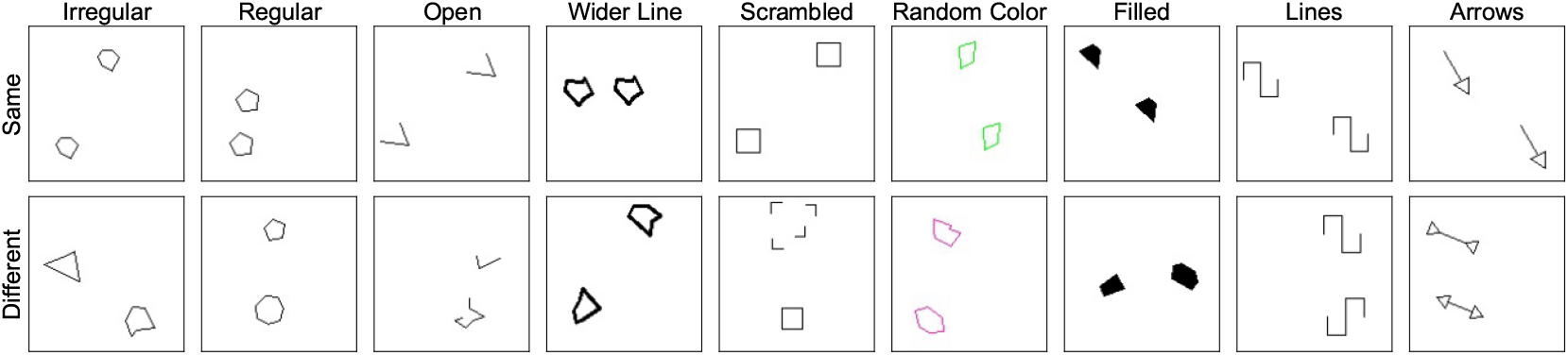
Examples of the “same” and “different” categories from our nine new versions of SVRT problem #1. See text for details.

Interestingly, Kim, Ricci, and Serre (2018) did find that a Siamese Network (Bromley, Guyon, LeCun, Säckinger, & Shah, 1993) that encoded the two shapes in two separate channels in order to simulate the effects of attentional selection and perceptual grouping, learned to classify images as “same” or “different” easily, leading the authors to conclude that object individuation is a key step in solving the same-different task. At the same time, they also argue that a full solution to the same-different problem requires a network to encode dynamic representations of relations rather than statically storing visual-relation templates in synaptic weights. That is, in their view, symbolic processes need to be implemented to fully solve the same-different task.

On the other hand, there are recent reports that the current state-of-the-art DCNNs can solve the same-different task. If this is indeed the case, it would be a striking example of standard networks solving a fundamental relational reasoning task without implementing any symbolic machinery. Funke et al. (2021) noted that Kim et al. (2018) only tested relatively small CNNs (up to 6 layers), and when they replicated the same-different experiments on the SVRT dataset using a `ResNet-50` (He, Zhang, Ren, & Sun, 2016) model (a network of 50 layers) the models were able to perform the task successfully. Funke et al. (2021) noted that the success does not necessarily imply DCNNs can solve all visual reasoning tasks, but they do highlight that standard feedforward processing DCNNs can solve the same-different task and that Kim et al.’s claim regarding the need for extra mechanisms for abstract visual reasoning is unwarranted.

Similarly, Messina, Amato, Carrara, Gennaro, and Falchi (2021) have shown that a range of recent DCNNs, specifically `ResNet, DenseNet` (Huang, Liu, Van Der Maaten, & Weinberger, 2017), and `CorNet-S` (Kubilius et al., 2018), can solve the same-different SVRT tasks, whereas they confirm that this is difficult for older `AlexNet` (Krizhevsky, Sutskever, & Hinton, 2012) and `VGG` (Liu & Deng, 2015) networks. The authors conclude: “We think that the development of the abstract and relational abilities of neural networks is an important leap towards achieving some interesting new tasks…”.

However, there is a fundamental problem with using success on the SVRT dataset as evidence that CNNs can support same-different relational reasoning. A key feature of relational reasoning is that it is reasoning based on relations between objects rather than any low-level visual details of the inputs. In the domain of visual reasoning, this entails that same-different discrimination should extend to novel images. The SVRT dataset does test models on novel images, but the test images are generated in the same way (i.e., the train and test datasets come from the same pixel-level distribution), and accordingly, it does not test the hypothesis that models have acquired the capacity to support relational reasoning on the same-different task.

## 2 Simulations

In the simulations below we test abstract same-different reasoning in several DCNNs models based on the `ResNet-50` architecture. The basic tenet of our simulations is that a model that has learned the abstract *same* and *different* relations should be able to recognize examples of these relations beyond its training set.

Our training and test data are based on problem #1 of the SVRT (see Fig. 1). In this problem images of two randomly generated shapes are classified as “same” if they are the same up to translation on the canvas and “different” otherwise. We created nine new datasets that followed the same abstract rule as problem #1 (see Fig. 2). However, each new dataset was generated through a distinct stochastic generative process (i.e., a different pixel-level distribution). Each dataset was defined as follows:

- In the Irregular dataset each shape was a irregular polygon. These polygons were generated by sampling a series of 1 to 7 points of a circumference of radio *π* (uniformly sampled from 1 to 40 pixels) around a randomly chosen center. After this, we added uniformly distributed random noise to each point and connected all of them with straight lines. In the “same” category both shapes were identical except for the position in the canvas. In the “different” category the initial points and point errors of the second polygon were re-sampled such that both shapes were different.
- In the Regular dataset each shape was a regular polygon. These polygon were generated in the same way as the Irregular dataset except that we did not add with random noise to the polygon points.
- The Open dataset was generated in the same way as the Irregular dataset except that the first and last vertices of each shape were not connected.
- The Wider Line dataset was generated in the same way as the Irregular dataset except that the line width was set to two pixels instead of one.
- The Scrambled dataset was was generated in the same way as the Regular dataset except that in the “different” category one the of the objects (scrambled) was generated by dividing the other object into sections and displacing them randomly around the center.
- The Random Color dataset was generated in the same way as the Irregular dataset except that for each image the line color was chosen randomly.
- The Filled dataset generated in the same way as the Irregular dataset except that the shapes were filled with black.
- In the Lines dataset each object corresponded to a line created by joining two open squares, one with the opening pointing downward and the other with the opening pointing upward, at the end of the opposite left/right sides (i.e., the right side downward-facing open square was joined to the left side of the upward-facing square or vice versa); in the “same” category the lines were identical whereas in the “different” category the second line was created by joining the open squares at the opposite left/right sides than the first line.
- In the Arrows dataset the objects were arrows consisting of one or two triangular head(s) and a line; the head(s) and the line were connected; in the “same” category the arrows were the same and in the “different” category the orientation of each head was inverted.

Note that among these nine different stimulus sets there are differences in the level of low-level similarity with the original SVRT data. In particular, the irregular, regular, and, to a lesser extent, the open datasets are more similar to the original data than the rest of the datasets. The code to generate these datasets, as well as to run all our simulations, can be found at https://github.com/GuillermoPuebla/same_different_paper.

### 2.1 Simulation 1

In Simulation 1 we performed the most basic and stringent test of abstract relational reasoning. We trained several models on the original problem #1 and then presented with 5600 images from each of the 10 stimulus test sets. That is, our testing conditions consisted on new images from the original training set (replicating Funke et al., 2021), and novel images from the other nine test datasets that were not seen during training. As noted above, a model that has learn the abstract *same* and *different* relations should generalize learning on the same-different task independently from the pixel-level similarity to original SVRT data.

In Simulation 1 we tested two sets of models based on the `ResNet-50` architecture. The first set consisted of four `ResNet-50` classifiers. All models consisted on a `ResNet-50` convolutional front end followed by a hidden layer with 1024 units with ReLU activation (see Fig. 3A). In Simulation 1 there was one output layer consisted of a single sigmoid unit which predicted the probability that the input image belonged to the category “same”. We pre-trained the models’ convolutional front end in either `ImageNet` (Deng et al., 2009) or TU-Berlin (Eitz, Hays, & Alexa, 2012), a dataset of human-generated sketches. Furthermore, we varied how we treated the output of the convolutional front end before passing it to the hidden layer. We either applied a global average pulling (GAP) operation to it, as Funke et al. (2021) did, or flatten it, as Messina et al. (2021) did^1^.

**Figure 3:**
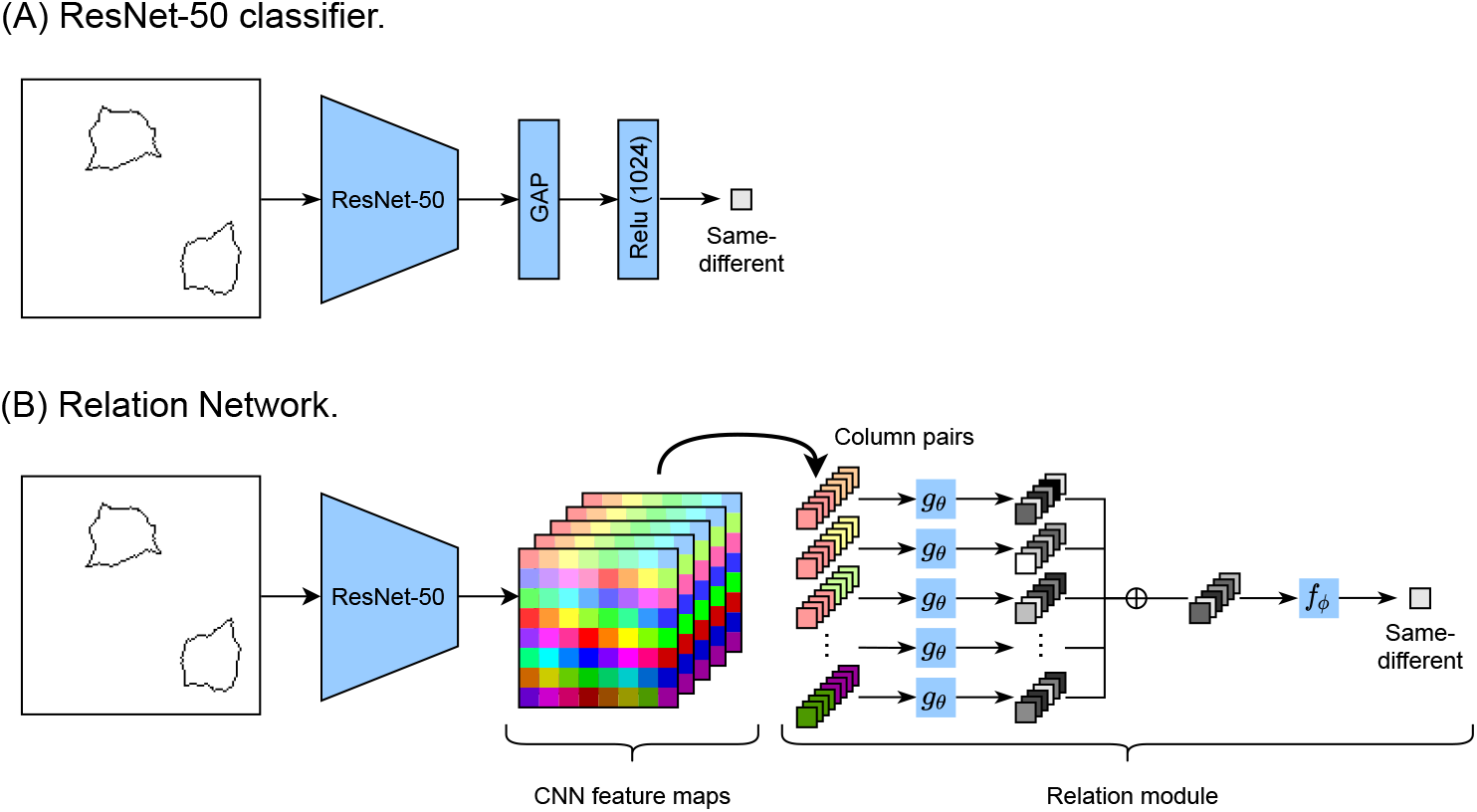
Models tested. (A) `Resnet-50` classifier. (B) Relation network.

The second set of models consisted of two variations of a Relation Network (Santoro et al., 2017). This architecture is especially relevant for the present study because it was explicitly designed to perform relational reasoning on the visual domain and it’s fully compatible with DCNNs. As ilustrated in Fig. 3B, a Relation Network consist of a convolutional front end, that outputs a series of filters, and a relation module. The relation module organizes the filter activations into columns that correspond to specific positions across filters (denoted by different colors in Fig. 3B), and generates all possible pairs of columns. All this pairs are processed by a single multilayer perceptron, *g_θ_*, yielding a vector per pair. These vectors are summed up and passed trough a second multilayer perceptron, *f_ϕ_*, that yields the final same/different prediction. Note that the feature columns inputs to the relation module do not necessarily represent objects or objects parts. Instead, they represent whatever is at their corresponding receptive fields (e.g., the background, a texture, or even multiple objects at the same time). We created two versions of the Relation Network^2^ by varying the filter inputs to the relation module. In the first version we used the output of the last convolutional layer of `ResNet-50` (pretrained on `ImageNet`), which consisted of 2048 4 × 4 filters. Because the original Relation Network of Santoro et al. used a CNN front end with filter outputs of size 8 × 8, in the second version we used the 1024 output filters of the last convolutional layer of `Resnet-50` with filter size 8 × 8.

Following the recommendations of (Mehrer, Spoerer, Kriegeskorte, & Kietzmann, 2020) that analyses of neural networks behaviour should be based on groups of network instances, we trained 10 instances of each model. We used the Adam optimizer (Kingma & Ba, 2014). Training proceeded in two stages. In the first stage, the pre-trained `ResNet-50` network was frozen while the rest of the network was trained with a learning rate of 0.0003. In the second stage, the complete model was trained with a learning rate of 0.0001. The training data consisted of the original data of the SVRT problem #1. In the first stage the model was trained on 28000 images for 5 epochs with a batches of 64 samples. In the second stage the model was trained on the same images for 10 epochs and with the same batch size.

Because same/different decisions were often performed on test datasets with different distributions than the training datasets, it is possible that there is a different optimal classification threshold for each test dataset. To account for this, we used the area under the ROC curve (AUC), which is a performance measure that takes into consideration all possible classification thresholds. AUC values range from 0.0 to 1.0, where 0.5 corresponds to chance-level responding and 1.0 corresponds to perfect classification. The AUC can be interpreted as the probability that a randomly sampled example of the positive category (“same”) will be assigned a higher predicted probability than a randomly sampled example of the negative category (“different”) (Hanley & McNeil, 1982). We interpreted the AUC values according the general guidelines of Hosmer, Lemeshow, and Sturdivant (2013, table 1).

**Table 1:** AUC interpretation criteria.

| AUC range | Category |
| --- | --- |
| $[0.9 - 1.0]$ | Outstanding |
| $[0.8 - 0.9)$ | Excellent |
| $[0.7 - 0.8)$ | Acceptable |
| $\leq 0.7$ | Poor |

#### 2.1.1 Results and Discussion

##### ResNet-50 Classifiers

As can be seen in Fig. 4, all models achieved outstanding performance in the Original test dataset. Furthermore, the models pre-trained on `ImageNet` performed better than the models pre-trained on TU-Berlin and the models with GAP performed better than the models that flattened the last convolutional layer’s output. Overall, the `ImageNet` & GAP model was the best performing model in the Original test dataset as well as, on average, in the nine new test datasets. Accordingly, the following analysis (as well as Simulations 2-4) will concentrate on it. The `ImageNet` & GAP model showed outstanding or excellent performance in the Irregular, Regular and Open datasets. As can be appreciated in Fig. 2, these datasets were the most featurally similar to the training data. On the other hand, the `ImageNet` & GAP model showed a poor performance on the Random Color, Filled, Lines, and Arrows datasets and a acceptable performance on the Wider Line and Scrambled datasets. In general, these results show that the degree of generalization on the same-different task depends heavily on the pixel-level similarity between the training data and the test data. This pattern of results is inconsistent with the models learning the abstract *same* and *different* relations.

**Figure 4:**
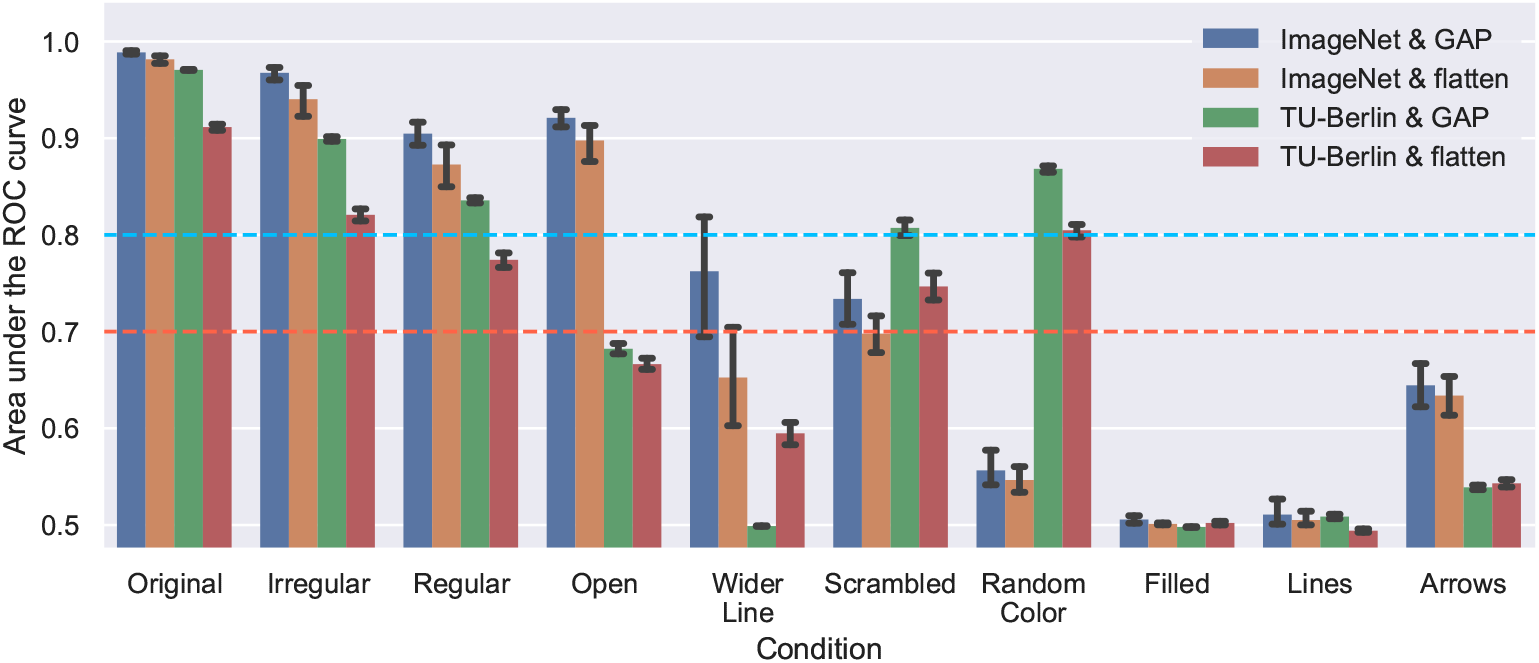
Mean AUC by ResNet-50 classifier and test dataset on Simulation 1. Error bars are 95% confidence intervals.

##### Relation Networks

Similarly to the `ResNet` classifiers, both model variations achieved outstanding performance in the Original test dataset (Fig. 5). Overall, the Relation network with 8 × 8 filter inputs was the best performing model in the Original test dataset as well as across the nine new test datasets (in many cases by a large margin). Accordingly, the following analysis (as well as Simulations 2-4) will concentrate on it. This model achieved excellent performance or above in the Irregular, Regular, Open, Wider Line, Scrambled and Random Color datasets. Performance was acceptable on the Filled dataset. Even though these results are better than the ones of the `ResNet` classifiers, the Relational network performed poorly on the Arrows and Lines datasets, which is inconsistent with an understating of the abstract relations *same* and *different*.

**Figure 5:**
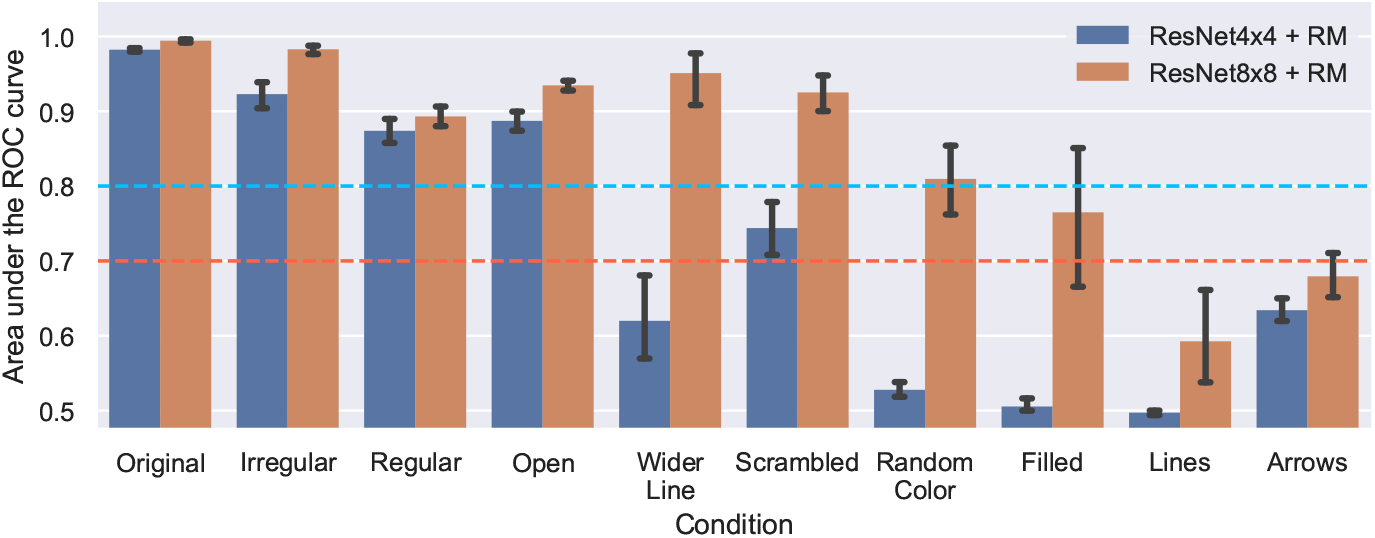
Mean AUC by Relation Network version and test dataset on Simulation 1. Error bars are 95% confidence intervals.

### 2.2 Simulation 2

One potential criticism to Simulation 1 is that the training data (line drawings of random shapes) wasn’t rich enough for the models to form a more complex representation of the “same” and “different” relations. Recall, however, that Messina et al. (2021) do interpret their results with the same training data as our Simulation 1 as supporting relational same-different reasoning in DCNNs and that Funke et al. (2021) interpret their results in terms of abstract visual reasoning. Nevertheless, we agree that the representations of the human visual system are based on rich stimuli and therefore is important to test what happens when the models have access to a richer training set. Therefore, in Simulations 2-4 we tested whether augmenting the training regime of the `ResNet-50`-based models would improve generalization on the same-different task to unseen stimuli. In Simulation 2, we did this by training the models on nine stimulus conditions consisting of images from the original SVRT data and all the new datasets except one. For each condition we trained 10 models with the same settings as in Simulation 1 except that the models were trained for 13 epochs instead of 10. We tested the models in the one stimulus set they were not trained on. For example, the models in the irregular stimulus condition were trained on the original data and all the new datasets except the irregular, in which they were tested on.

#### 2.2.1 Results and Discussion

As can be seen in Fig. 6, the `ResNet-50` classifier and the Relation Network performed similarly across the test datasets. A notable exception was the Random Color datatset, were the Relation Network achieved acceptable performance (borderline excellent) whereas the `ResNet-50` classifier showed poor performance. Overall, these results show that augmenting the training regime directly on the same-different task improved performance on untrained datasets for both models. However, this benefit seems more related to the pixel-level similarity of the augmented dataset with the testing data than to the shared relational structure of the problem among conditions, as illustrated by the results on the Lines dataset, where both models responded at chance levels. If the data augmentation helped the models to learn the abstract *same* and *different* relations the generalization to unseen data would have spread to all datasets.

**Figure 6:**
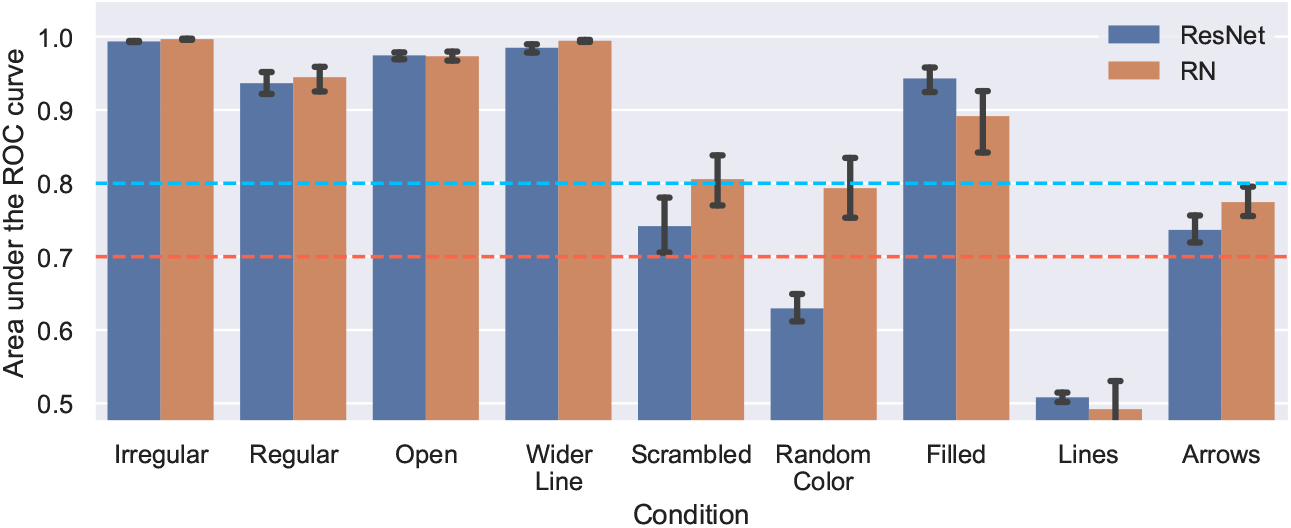
Mean AUC by test dataset and model on Simulation 2. Error bars are 95% confidence intervals.

### 2.3 Simulation 3

In Simulation 2 we augmented the models’ experience of the different conditions by training on the same-different task directly. A potential criticism to this strategy is that it does not give the models any experience with the specific condition they are tested on. In contrast, in Simulation 3 we augmented the models’ experience on all the datasets through multi-task learning. In deep learning research multi-task learning has long been used as technique to improve generalization (for a review, see Ruder, 2017). In this simulation the models were trained on two tasks. The first was the same-different task as in the previous simulations. The second was a relative position task. This consisted in classifying whether the lower object in the image was to the right of the upper object (category 1) or to the left (category 0).

To accomplish this on the `ResNet-50` classifier we added a second output layer with a single sigmoid unit. We trained 10 instances with images from the “same” and “different” categories from all conditions. However, we only allowed the models to learn to classify images from the original SVRT data as “same” or “different”, whereas the models learned to classify all presented images into their corresponding relative position category. To accomplish this, during training we used the following composed loss function:

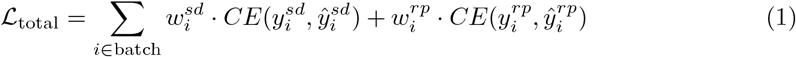

were *CE*(*y, ŷ*) is the cross-entropy loss between the label *y* and the prediction *ŷ*, and *w^sd^* and *w^rp^* are the weights for the same-different loss and the relative position loss, respectively. During training, *w^rp^* was set to 1 for all images. In contrast, when the model received images from the original SVRT data we set *w^sd^* to 1, otherwise it was set to 0. During testing, we presented the models with images of each problem version and recorded the models’ same-different and relative position AUC. All other training and testing parameters were the same as in Simulation 1 except that we trained the models for 13 epochs rather than 10.

For the Relation Network, we added a question layer^3^ and a second output layer with a single sigmoid unit. The Relation Network concatenates this question to all the column pairs, making the computation performed by *g_θ_* question-dependent. To train on the same-different task we created a vector, [1 0]^*T*^, representing the same-different question and another vector, [0 1]^*T*^, representing the relative position question. When the input to the question layer was the same-different vector the target for the relative position output was always 0, and when the input to the question layer was the relative position vector the target for the same-different output was always 0. We trained 10 instances of the Relation Network on two Original datasets, one with the input to the question layer corresponding to the same-different vector, and one with question input corresponding to the relative position vector. For all other datasets we set the question inputs to the relative position vector.

#### 2.3.1 Results and Discussion

As can be seen in Fig. 7, both models achieved perfect performance in the relative position task in all test datasets. On the other hand, in the same-different task the results were more varied. At least one of the models showed a excellent or outstanding performance in the majority of the test datasets except for Filled, Lines and Arrows. Strikingly, overall the `ResNet-50` classifier performed better than the Relation Network. Notably, both models performed poorly on the Lines and Arrows datasets despite having seen images from those datasets on the secondary relative position task. These results show that augmenting the model’s experience by training on a auxiliary task can enhance generalization on a target task for unfamiliar samples. However, the auxiliary task did not enhance generalization evenly across datasets, which suggest that the auxiliary task is not helping the model to form an abstract representation of the relations *same* and *different*.

**Figure 7:**
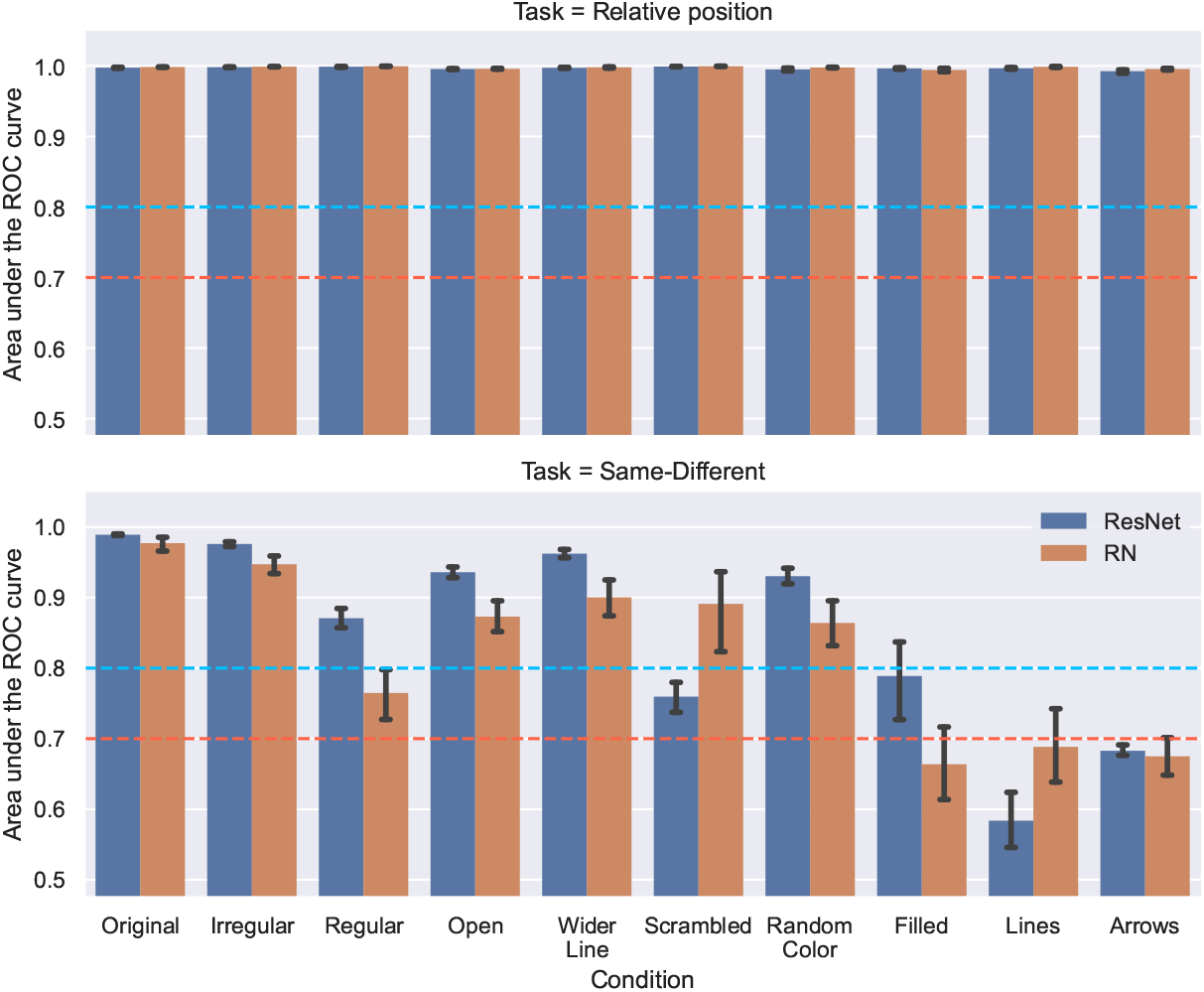
Mean AUC by test dataset, model and task on Simulation 3. Error bars are 95% confidence intervals.

### 2.4 Simulation 4

In Simulation 4 we combined the approaches taken in Simulations 2 and 3 in order to provide the models with the maximum amount of information to generalize the same-different task to the unseen conditions. As in Simulation 3, we trained the models in both the same-different and the relative position tasks. Furthermore, as in Simulation 2, for the same-different task we trained on all the stimulus conditions except one. For each of these 9 conditions we trained 10 instances of the `ResNet-50` classifier and the Relation Network and tested them on the stimulus set that was not trained on. We trained the `ResNet-50` classifier with loss (1), this time setting *w^sd^* to 1 for all datasets except the one tested on. All other training parameters were the same as in Simulation 3 for both models.

#### 2.4.1 Results and Discussion

As can be seen in Fig. 8, for the relative position task both models achieved perfect performance in all test datasets, just as in Simulation 3. Performance in the same-different task was better than in Simulation 3, with both models achieving excellent or outstanding performance in all test datasets except Scrambled, Lines and Arrows. Just as in Simulation 3, overall the `ResNet-50` classifier performed better than the Relation Network. Furthermore, despite being trained trained on all datasets on the relative position task and on all *other* datasets (i.e., except on the dataset tested) on the same-different task, the the `ResNet-50` classifier and the Relation Network only achieved an acceptable level of performance on the Arrows dataset and performed poorly on the Lines dataset. Overall, this pattern of results is not consistent with the models learning the abstract relational concepts *same* and *different*.

**Figure 8:**
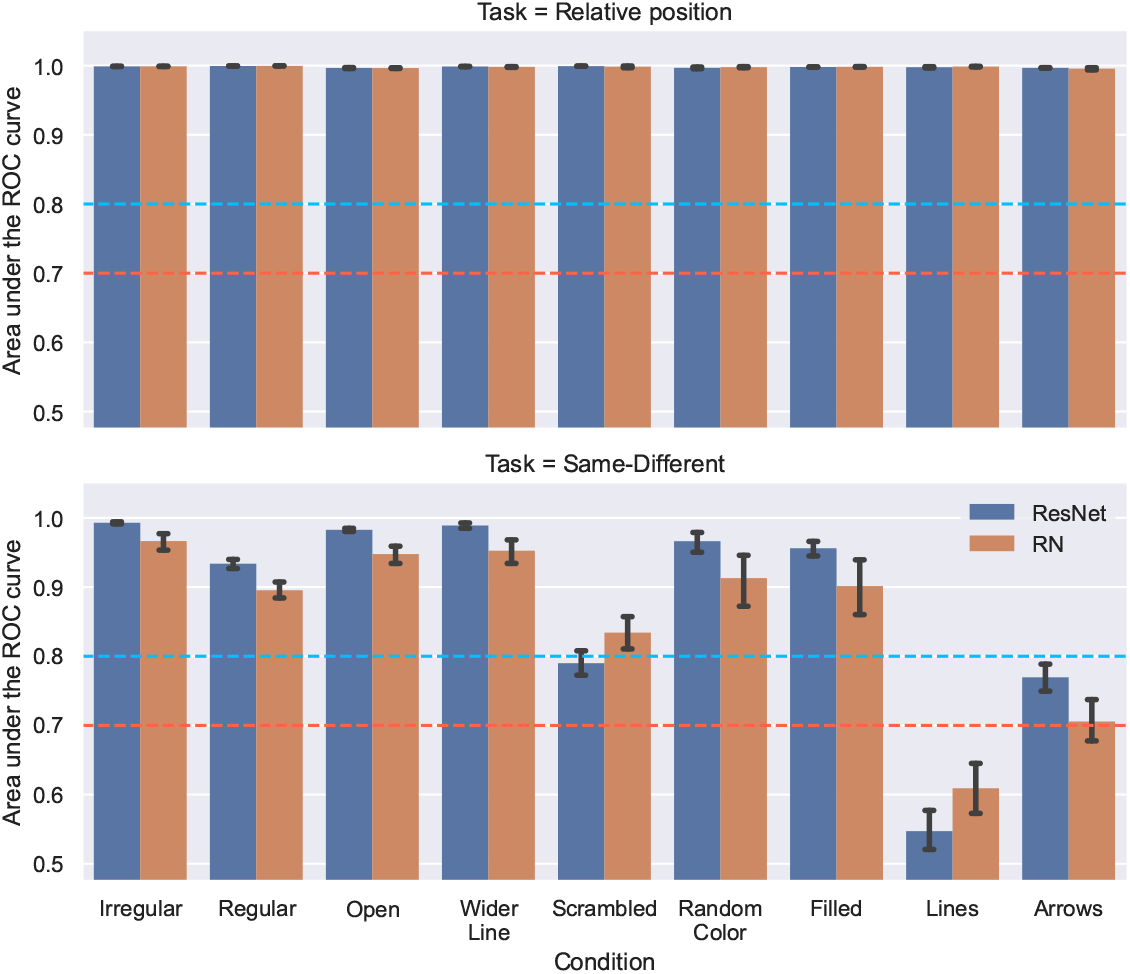
Mean AUC by test dataset, model and task on Simulation 4. Error bars are 95% confidence intervals.

### 2.5 Simulation 5

Simulations 1 to 4 showed that augmenting the model’s training regime, either directly by training on the same-different task with data from several datasets or indirectly by multitask learning, improved generalization in both the `ResNet-50` classifier and the Relational Network. As described previously, however, the benefits of both strategies did not spread to all test datasets equally, which raises the question of how much does this benefit extends to unseen examples of the “same” and “different” categories beyond what would be expected from pixel-level similarity. To test the extent of the generalization benefits of data augmentation, in Simulation 5 we trained the `ResNet-50` classifier and the Relational Network on all previous datasets on both tasks and tested them in four new test datasets (see Fig. 9). These datasets followed the same rule as SVRT problem #1, but differed from the previous conditions at the pixel-level. They were defined as follows:

- In the Rectangles dataset each shape was a rectangle. In the “same” category both shapes were identical whereas in the “different” category either the width or the height of one of the rectangles was different. Both sides had lengths between 16 and 64 pixels and the minimum difference between the critical sides on the “different” category was 4 pixels.
- In the Straight Lines dataset each shape was a straight line with a tilt of 0°, 45°, 90° or 135°. In the “same” category both lines were identical. In the “different” category one line was longer than the other. The lines had a length between 16 and 64 pixels and minimum difference in length was 4 pixels. This differences were uniformly distributed across examples of the “different” category.
- In the Connected Squares dataset each shape was a pair of connected squares. These shapes were generated by adding an horizontal line to the shaped of the Lines condition (compare the 3rd column of Fig 9 with the 8th column of Fig. 2). In the “same” category both shapes were identical. In the “different” category the corner at which both squares were connected was the opposite.
- In the Connected Circles dataset each shape was a pair of connected circles where one was in top of the other. One the circles was bigger than the other. In the “same” category both shapes were identical. In the “different” category the circles at the top and bottom were swapped.

**Figure 9:**
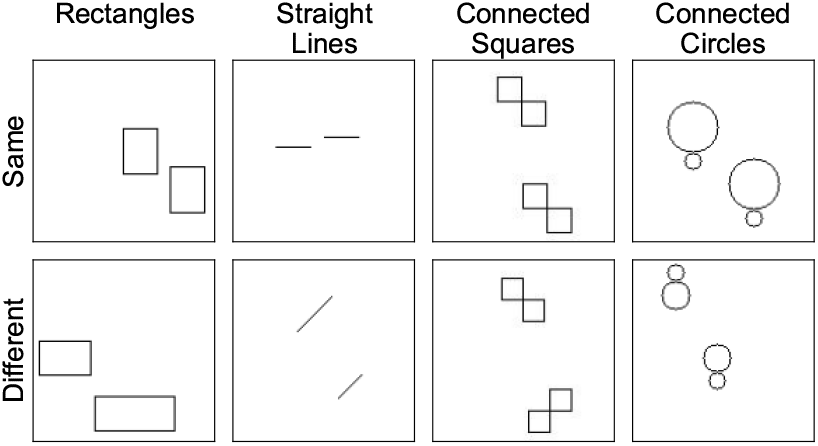
Examples of the “same” and “different” categories from the new testing conditions of Simulation 5.

To train the models we used the same settings as the previous simulations, except that we trained the 10 model instances for 19 epochs.

#### 2.5.1 Results and Discussion

As can be seen in Fig. 10, both models achieved ceiling performance on the relative position task in all the new test datasets. In contrast, both models only achieved acceptable performance on the Rectangles dataset and poor performance on the Straight Lines, Connected Squares and Connected Circles datasets. This marked difference in task generalization is consistent with previous results by Kim et al. (2018; see also Vaishnav et al., 2021), who reported that visual reasoning tasks that involve same-different judgments are harder for CNNs than other spacial reasoning tasks.

**Figure 10:**
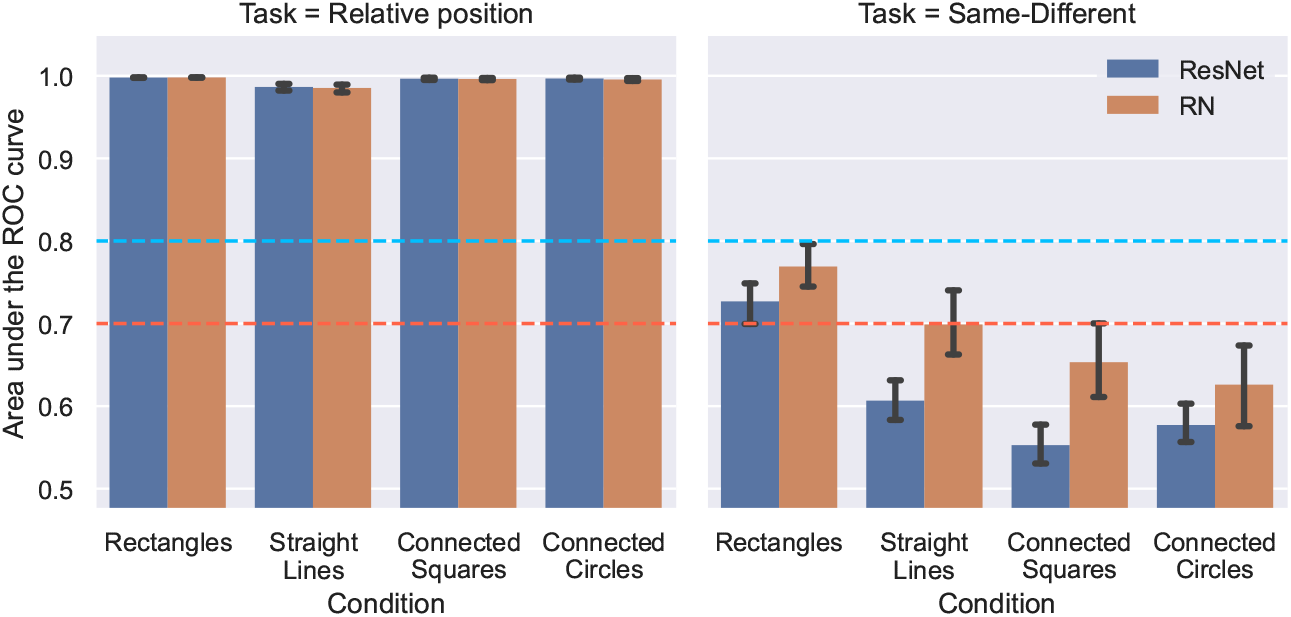
Mean AUC by test dataset, model and task on Simulation 5. Error bars are 95% confidence intervals.

In order to gain further understanding of both models’ behaviour in this simulation, for each dataset/model combination we picked the model instance that produced the median AUC score and used these instances to plot the distribution of predicted probabilities for each new test dataset for both models (Fig. 11)^4^. For each model/dataset combination we also calculated the optimal threshold that maximized the true positive rate and the true negative rate, through Youden’s J index (Youden, 1950), and superimposed on the distributions. As can be seen in Fig. 11, overall the distributions of predicted probabilities for the “same” and “different” categories showed a high degree of overlap. For both models the overlap was higher in the Connected Squared and Connected Circles datasets.

**Figure 11:**
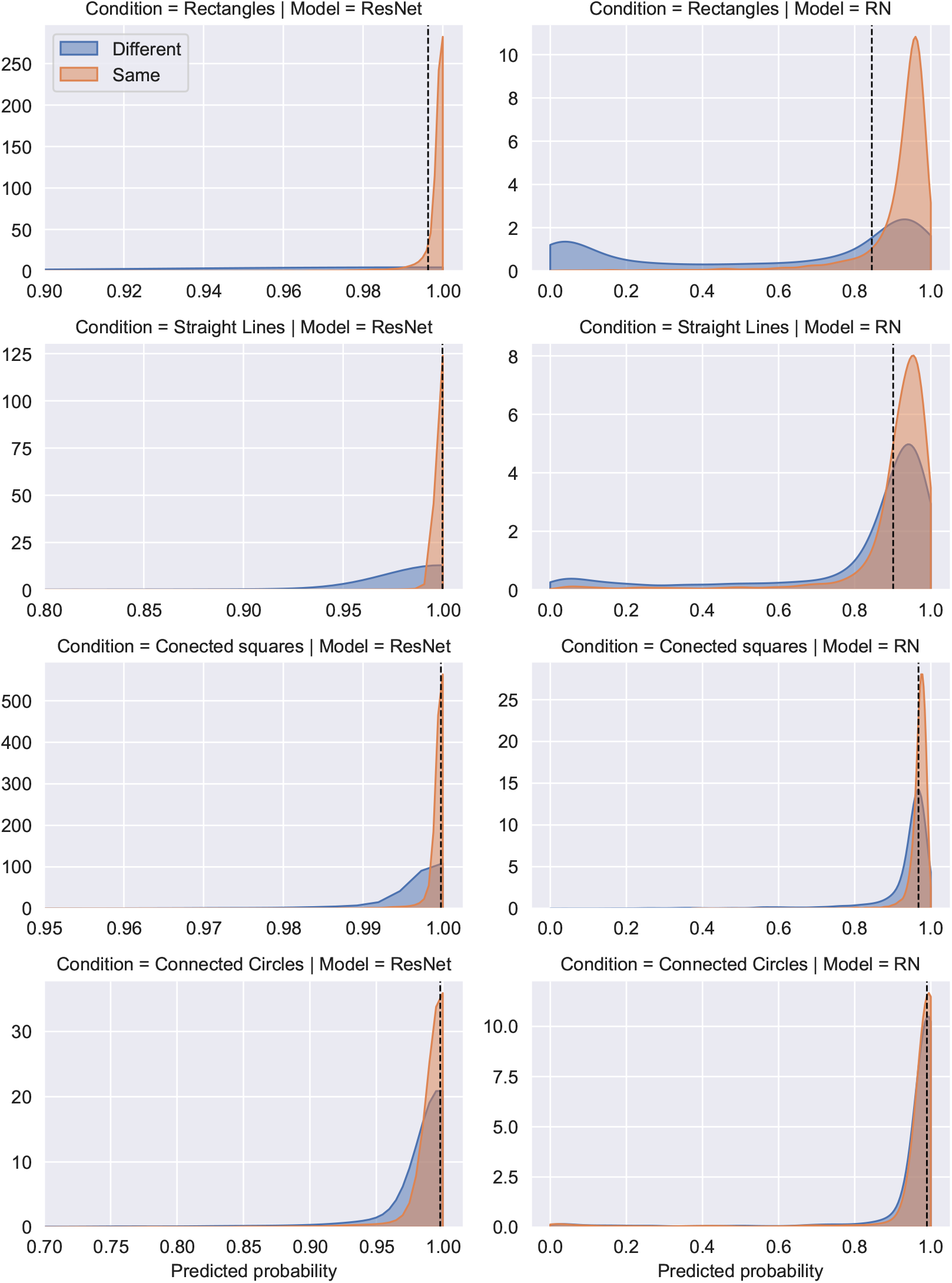
Gaussian kernel density estimates for new test datasets on Simulation 5. Dashed lines represent optimal thresholds each model/dataset combination. See text for details.

For the `ResNet-50` classifier the optimal threshold was very close to 1.0 in all datasets. In the Rectangles dataset this meant that the model would classify most examples of the “same” category correctly, as well as most cases of the “different” category, although the distribution for the “different” category was very flat, which means that the model was as likely to assign a low predicted probability to this examples as a high one. In contrast, the high degree of overlap of the distributions on the rest of the datasets made the model to classify the majority of the examples of both categories incorrectly even when using the optimal threshold.

For the Relation Network the overlap of the distributions in the Rectangles dataset was less pronounced, which made the optimal threshold being able to classify correctly most of the examples of the “same” category, and large proportion of the examples of the “different” category correctly, although a significant proportion of cases still remains misclassified. In all other conditions the higher overlap between the distributions made the optimal threshold to classify incorrectly a larger proportion of the examples of the “same” and “different” categories.

Overall, the results of Simulation 5 show that generalization of the same-different task was highly constrained by the pixel-level similarity of the testing examples to the training data for both, the `ResNet-50` classifier and the Relation Network, exactly the opposite of one would expect if these models had learned the abstract relations *same* and *different*.

### 2.6 Simulation 6

Simulations 1 to 5 showed that DCNNs do not learn the abstract *same* and *different* relations. What is needed to accomplish this? As discussed in the introduction, Kim et al. (2018; see also Ricci et al., 2021) have argued that the Siamese Networks (Bromley et al., 1993), a model that simulates the effects of attentional selection and perceptual grouping by encoding the two shapes of the same-different examples in two separate channels, was able to solve the same-different task easily.

However, there are reasons to doubt that separating the shapes of the same-different examples into different channels is all that is needed for a DCNN to learn an abstract representation of the relations *same* and *different*. First, there has been no tests of the generalization capabilities of the Siamese Network on the kind challenging stimuli we have used on Simulations 1 to 5. Second, recently Webb, Sinha, and Cohen (2021) have shown, using a custom dataset, that a recurrent version of the Siamese Network failed on the same-different task on examples not seen during training. In contrast, an Emergent Symbol Binding Network (ESBN) that used a two-stream external memory, where one stream was populated with CNN embeddings and the other was populated by the hidden state vectors of a recurrent neural network, was able to generalize same-different discrimination much better. Webb et al. (2021) argue that this separation, coupled with a similarity-based mechanism to bind the representation of the hidden state vectors with the representations of the objects, allowed the ESBN to represent the same-different task in terms of abstract variables.

In order to investigate the role of channel separation in same-different generalization without implementing extra symbolic machinery, in Simulation 6 we replicated Simulations 1 and 5 with a Siamese Network. Our model (Fig. 12), uses the same `Resnet-50` in both channels (i.e., they share weights) to produce two vectors through a GAP operation, that are concatenated and passed to two hidden layers of ReLU units which lead to a single same/different output unit. To train and test the model, we made versions of all our new datasets with the objects separated into two images. In Simulation 6a we trained the model on the Irregular dataset and tested it in all other datasets (as in Simulation 1). In Simulation 6b we trained on the same-different task in all the datasets of Simulation 6a and tested on the Rectangles, Straight Lines, Connected Squares and Connected Circles conditions (as in Simulation 5). We used the same training settings as the previous simulations, except that we trained the models for 5 epochs (at which point all 10 model instances achieved ceiling performance on the training data).

**Figure 12:**
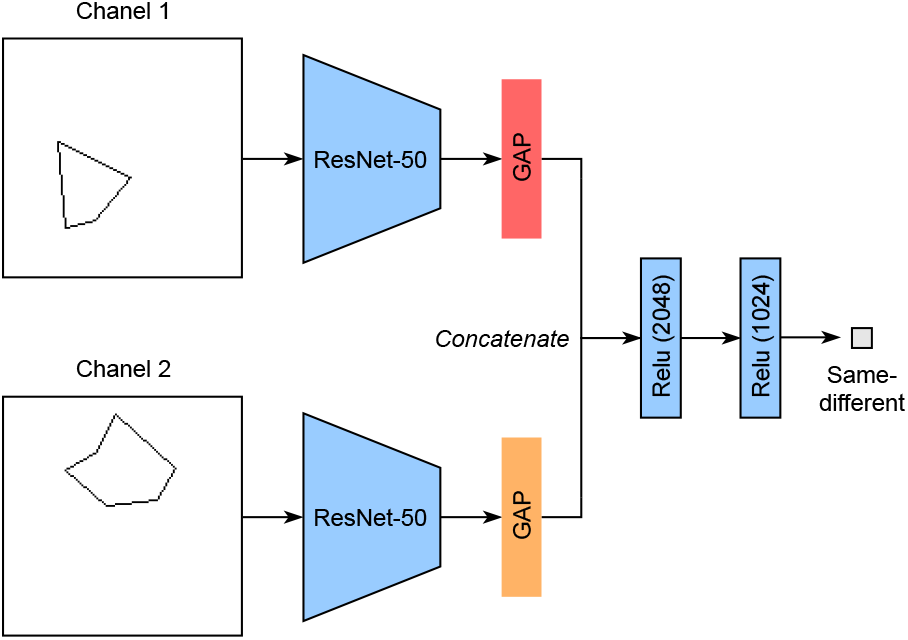
Siamese Network.

#### 2.6.1 Results and Discussion

As can be seen in the first panel of Fig. 13, Simulation 6a shows that training on the separated channels version of the Regular dataset did not produce perfect generalization to the untrained datasets, with the model achieving acceptable performance on the Arrows dataset and poor performance on the Scrambled and Lines datasets. Furthermore, Simulation 6b (Fig. 13 second panel) shows that training in all previous datasets did not produce perfect same-different generalization either. It is worth noting, however, that Simulation 6b produced better results than Simulation 5, with performance on the Connected squares dataset going from poor to excellent.

**Figure 13:**
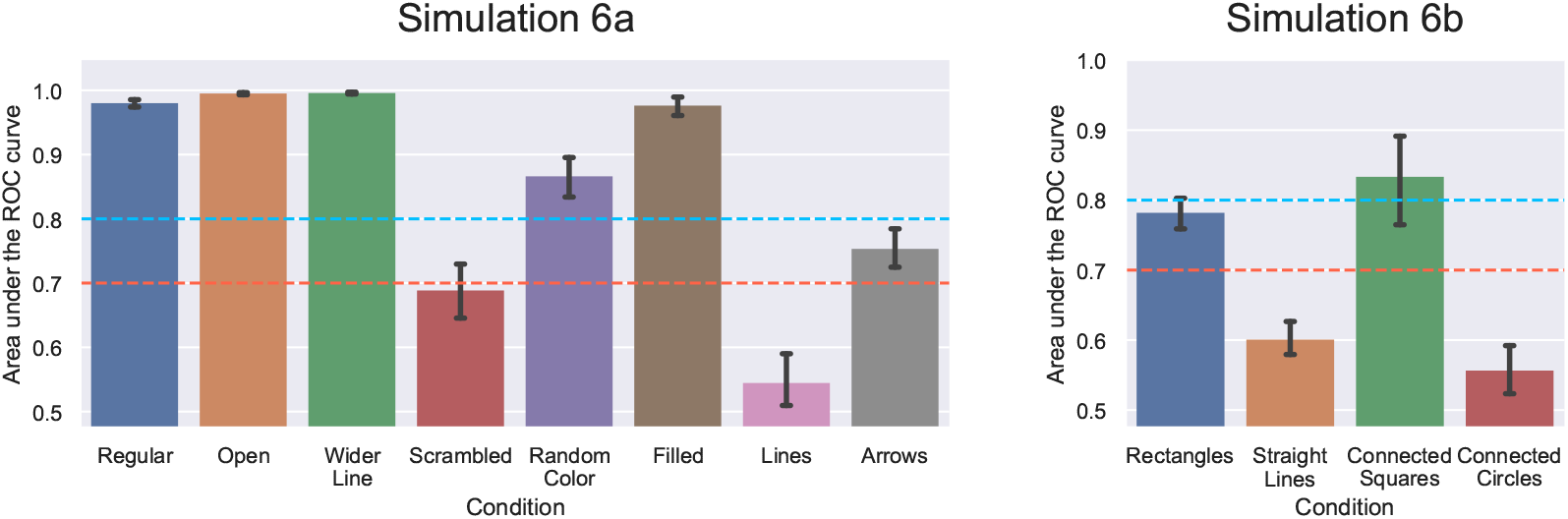
Mean AUC by test dataset on Simulations 6a and 6b. Error bars are 95% confidence intervals.

Overall, these results show that that using separated channels for the two objects of each example did not produce perfect generalization on the same-different task. These results imply that object individuation is not a sufficient condition for learning the relations *same* and *different* on DCNNs. Most likely, besides object individuation it is necessary to separate the content of the representations of the objects participating in the relation from the content of the relational roles that form the abstract relations *same* and *different*, as postulated by part-based theories of object recognition (Biederman, 1987; Hummel & Biederman, 1992) and implemented in the ESBN model of Webb et al. (2021).

## 3 General Discussion

In six simulations we tested whether DCCNs were able to learn the abstract *same* and *different* relations when trained on the same-different task. Across simulations we found that, instead of forming an abstract representation of this task that generalizes beyond the training distribution, DCCNs were unable to generalize to new test images that shared the same underlying relations as the training data but were dissimilar at the pixel level. This was the case even when we augmented DCCNs’ experience with new stimulus sets that instantiated the same-different task with several kinds of objects (Simulations 2, 4 and 5), and when we used multi-task learning to give them experience with the very same images in a different task (Simulations 3, 4 and 5). Furthermore, in Simulation 6 we showed that separating the two objects of the same-different images into separated channels was not enough to enable DCNNs to learn the abstract notions of *same* and *different*, even when trained in a rich regime with data from several datasets.

These results shed new light into the discussion of whether is necessary to invoke extra, symbolic mechanism to solve the same-different task. If by “solving” the same-different task one means generalizing from one set of images to another set of images that share the same pixel-level distribution, as implicitly assumed by Messina et al. (2021) and Funke et al. (2021) and implemented in the SVRT dataset, it is perfectly reasonable to say that DCNNs are able to solve this task. This, by itself, is an interesting problem from a machine learning point of view, because simpler machine learning models tested previously could not solve this kind of task. However, if by “solving” the same-different task one means to learn a representation of the *same* and *different* relations that support generalization beyond pixel-level similarity (as in humans and chimpanzees), our results suggest that DCCNs are just not up to the task and that some sort of symbolic machinery may be necessary.

Consistent with this conclusion, the Relation Network used in Experiments 1-5 did not fair much better than standard CNNs classifiers when the training and test images were markedly different at the pixel level. Importantly, the Relation Network is claimed to support relational reasoning without implementing symbolic computations. Our results show, however, that exhaustively comparing feature columns from all locations in the output filters of a DCNN does not support out-of-distribution generalization of same-different judgements, as one would expect from a model that learned a relational representation of the same-different task. Besides failing to learn what could be considered the simplest possible visual relations, the number of parameters of the Relation Network grows combinatorially with the filter size of the DCNN output, which makes this model much more inefficient during training than standard CNNs classifier and brings into question the scalability of this approach^5^.

Similarly, the Siamese Network that does not implement any symbolic computations failed to support same/different judgments when training and test images were from different pixel-level distributions. Clearly, object individuation is a necessary step in the process of comparing objects, but our findings highlight that hard-wiring this information in a DCNN is not sufficient in to solve the same-different task. As many have suggested (e.g., Webb et al., 2021; Hummel & Biederman, 1992, for a review see Greff, van Steenkiste, & Schmidhuber, 2020), for a neural network to achieve effective relational generalization, mechanisms to represent objects and relational roles independently and binding them together dynamically might be necessary. The results of Webb et al. (2021) with the ESBN model could be seen as supporting this hypothesis. Note, however, that the dataset of Webb et al. (2021) was composed by combining 100 32 × 32 grayscale images of simple Unicode characters, which raises the question of whether the ESBN model would show the same degree of generalization with more complex stimuli like ours. We think that this is an interesting possible extension of the current research.

More generally, as Stabinger et al. (2020) note, models that work with separated channels for different objects assume one of the most important steps of the processing of visual relations. A satisfactory solution to the object individuation problem in neural networks should be able to extract the critical objects to compare from the image automatically. As far as we are aware, there is not a DCNN capable of this feat.

In conclusion, our results show learning same-different relations is beyond the current capabilities of DCNNs. Fundamental work on mechanisms for object individuation and dynamic binding seems necessary for neural networks achieve this hallmark of intelligence.

## Appendix

We benchmarked the Relation Network used on Simulation 1 on the Sort-of-CLEVR dataset (Santoro et al., 2017, Fig. 14). This dataset consists of images with 6 objects. Each object has a unique color (red, green, blue, orange, gray, or yellow) and it has a square or a circular shape. Each image has 20 associated questions, 10 of which are non-relational and 10 are relational. The non-relational questions ask for (a) the shape of an object, (b) the horizontal location of an object (left or right) or (c) the vertical location of an object (upside or downside). These questions are considered non-relational because to answer them a model needs to focus only on a single object. On the other hand, the relational questions require the models to consider relations between the objects in the image. These questions ask for (a) the shape of the object which is closest to certain object, (b) the shape of the object which is furthest away from certain object, and (c) the number of objects that have the same shape as certain object. The dataset consisted of 10000 randomly generated (image, questions, answers) triplets, of which 200 were withhold for testing. In order to benchmark the `ResNet-50` based Relation Network, we made images of size 128 × 128 instead of 75 × 75 as in the original dataset.

**Figure 14:**
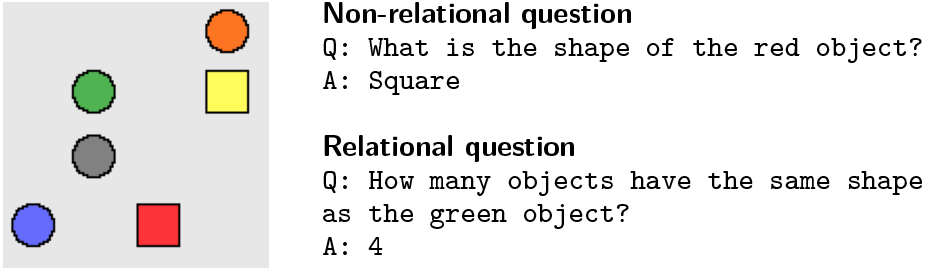
Example image, questions and anwsers from the Sort-of-CLEVR dataset.

We benchmarked three models. The first two were the same models tested in Santoro et al. (2017): a four later CNN front end with a MLP classifier (CNN+MLP) and the original Relation Network. The third one was the Relation Network with 8×8 inputs from `Resnet-50`. All models were trained with the Adam optimizer with a learning rate of 0.00025 for the first two models and 0.0001 for the last. As can be seen in Fig. 15, both versions of the Relation Network achieved high levels of performance in the non-relational and relational questions. The CNN+MLP model performed comparatively worse in both types of questions. In contrast to the results of Santoro et al. (2017), the CNN+MLP model performed better in the relational questions than in the non-relational ones. It is worth noting that the Sort-of-CLEVR dataset does not include a set of withhold objects, which prevents to perform the kind of relational generalization tests that we carry on the main article. In fact, when Kim et al. (2018) tested the Relation Network on a same-different dataset with withhold color/shape combinations the model performed at chance.

**Figure 15:**
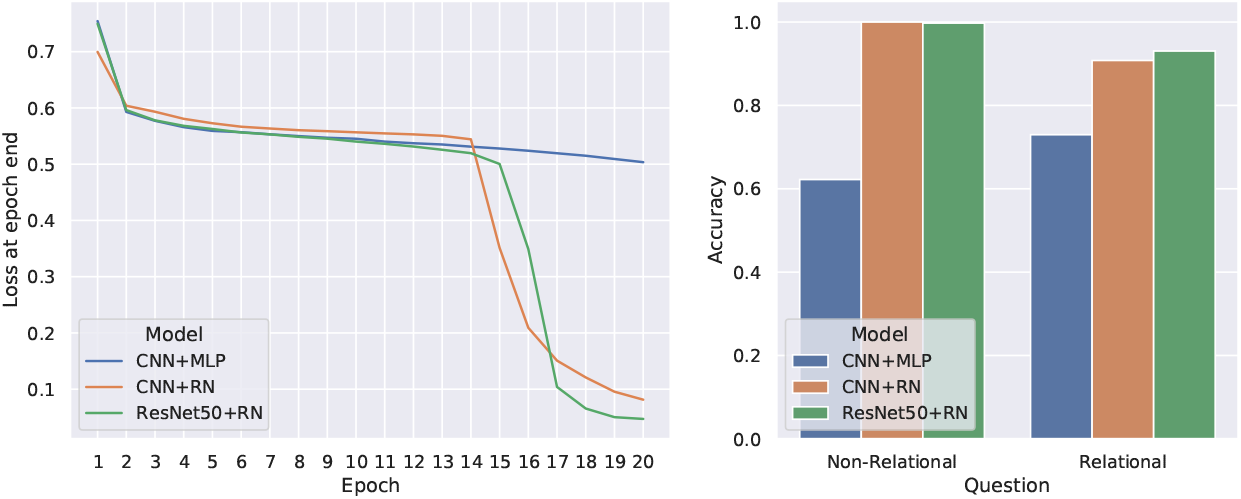
Sort-of-CLEVR benchmark. Left: training loss. Right: test accuracy on non-relational and relational questions.

## Footnotes

^1^We also made models that had the pre-trained convolutional front end frozen and only the classifier was trainable. Those models achieved similar results to the ones presented on Simulation 1. However, they were not well-suited for the data augmentation and multi-task learning techniques used on Simulations 2-4, so we don’t consider them further.

^2^To validate our implementation of the Relation Network we replicated the main results of Santoro et al. (2017) with the Sort-of-CLEVR dataset (see, Appendix).

^3^The question layer was part of the original implementation of the Relation Network by Santoro et al. (2017) but was unnecessary in the previous simulations because the question was constant (same or different).

^4^ We used the kernel density estimate function of Seaborn (Waskom, 2021). We customized the y-axis limits for each dataset on the `ResNet-50` classifier column for better readability

^5^ In our simulations this made training the Relational Network several times slower than training a the `ResNet-50` classifier using a GPU.

